# Neuro-immune modulation of cholinergic signaling in an addiction vulnerability trait

**DOI:** 10.1101/2022.12.05.519158

**Authors:** Hanna Carmon, Evan C. Haley, Vinay Parikh, Natalie C. Tronson, Martin Sarter

**Author notes:** **Author contributions:** HC determined the rats’ phenotype and measured cytokine levels in brain and spleen, determined the effects of LPS on cytokine concentrations, and prepared brain tissues for measurement of choline transporter concentrations and modifications. ECH and VP determined subcellular choline transporter concentrations, ubiquitinylation levels, and the effects of LPS. VP, NCT and MS designed the experiments. NCT and MS wrote the original draft of the paper. All authors contributed to the statistical analyses of the data and to the editing of the final manuscript. **Corresponding author:** Martin Sarter, University of Michigan, Dept. of Psychology, 530 Church Street, Ann Arbor, MI 48109.

## Abstract

Sign-tracking (ST) describes the propensity to approach and contact a Pavlovian reward cue. By contrast, goal-trackers (GTs) respond to such a cue by retrieving the reward. These behaviors index the presence of opponent cognitive-motivational traits, with STs exhibiting attentional control deficits, behavior dominated by incentive motivational processes, and vulnerability for addictive drug taking. Attentional control deficits in STs were previously attributed to attenuated cholinergic signaling, resulting from deficient translocation of intracellular choline transporters (CHTs) into synaptosomal plasma membrane. Here we investigated a post-translational modification of CHTs – poly-ubiquitination - and tested the hypothesis that elevated cytokine signaling in STs contributes to CHT modification. We demonstrated that intracellular CHTs, but not plasma membrane CHTs, are highly ubiquitinated in male and female sign-tracking rats when compared with GTs. Moreover, levels of cytokines measured in cortex and striatum, but not spleen, were higher in STs than in GTs. Activation of the innate immune system by systemic administration of the bacterial endotoxin lipopolysaccharide (LPS) elevated ubiquitinated CHT levels in cortex and striatum of GTs only, suggesting ceiling effects in STs. In spleen, LPS increased levels of most cytokines in both phenotypes. In cortex, LPS particularly robustly increased levels of the chemokines CCL2 and CXCL10. Phenotype-specific increases were restricted to GTs, again suggesting ceiling effects in STs. These results indicate that interactions between elevated brain immune modulator signaling and CHT regulation are essential components of the neuronal underpinnings of the addiction vulnerability trait indexed by sign-tracking.

**Significance Statement:** Sign-tracking rats (STs) have emerged as a fruitful animal model for determining the neuro-behavioral foundations, including the cholinergic-attentional control deficits, which heighten the risk for the manifestation of addictive behaviors. The results from the present experiments suggest that, in STs, elevated levels of neuro-immune signaling interact with a post-translational modification of choline transporters that yields a relatively low transporter capacity. Furthermore, the effects of activation of the innate immune system, by administrating the endotoxin lipopolysaccharide, mechanistically supported such an interaction. The results extend research on the neuronal vulnerabilities for addictive behaviors to the role of neuro-immune signaling and suggests a new role of neuro-immune modulators in influencing complex cognitive-motivational traits.

## Introduction

Repeated administration of addictive drugs *per se* does not reliably predict the manifestation of substance use disorder. Rather, neuro-behavioral vulnerabilities, such as fronto-cortico-striatal circuit abnormalities, impulsivity, and attentional control deficits, have been proposed to interact with the psychobiological effects of addictive drugs to foster compulsive drug taking (e.g., Mueller et al., 2020; Lamontagne and Olmstead, 2019; Marhe et al., 2013; Field and Cox, 2008; Tomasi et al., 2007; Ersche et al., 2012; Ersche et al., 2011; Makris et al., 2008; Redzepagic and Ladas, 2023; Anselme and Robinson, 2020). Sign-tracking indicates the presence of such a vulnerability trait. Using a Pavlovian Conditioning Approach (PCA) test, sign-trackers (STs) approach and operate a Pavlovian reward cue (typically a lever). Their counterparts, the goal-trackers (GTs), also learn about the predictive significance of the cue but they do not approach it. Sign-tracking has been interpreted as reflecting a propensity for attributing incentive salience to Pavlovian cues (e.g., Meyer et al., 2012; Flagel et al., 2011; Saunders and Robinson, 2011, 2010; Robinson and Berridge, 1993; Schad et al., 2020).

Attentional control deficits, including a bias for cue-driven or bottom-up attention, have long been considered an essential component of addiction vulnerability traits (Franken, 2003; Tomasi et al., 2007; Field and Cox, 2008). Approaching a non-instrumental reward cue in part may reflect the attentional “magnetism” of such cues for STs. In contrast, the behavior of GTs during PCA testing is directed solely toward the food port, perhaps reflecting goal-directed attention and model-based processing (Clark et al., 2012). Consistent with this view, STs perform relatively poorly when tested in a sustained attention task and in situations taxing the attentional supervision of complex movements. These phenotype characteristics have been attributed to a limited capacity of basal forebrain-cortical cholinergic projections to support such behaviors (Paolone et al., 2013; Pitchers et al., 2018; Pitchers et al., 2017b; Pitchers et al., 2017a; Phillips and Sarter, 2020; Kucinski et al., 2019; Kucinski et al., 2018; Kucinski et al., 2022).

A cellular mechanism responsible for such limited cholinergic capacity in STs has previously been described. In STs, the translocation of neuronal choline transporters (CHTs) from intracellular domains into synaptosomal plasma membrane was found to be deficient, thereby failing to support increases in synaptosomal choline import, acetylcholine (ACh) synthesis and release (Koshy Cherian et al., 2017).

We first asked whether a post-translational modification of CHTs, poly-ubiquitination, is present in STs and thereby may contribute to their CHT trafficking deficit. Ubiquitination can occur in diverse ways, including conjugation to just one residue in its substrate (mono-ubiquitination) or multiple times to this residue, with the initial ubiquitin serving as the target for additional ubiquitination (poly-ubiquitination). Beyond the classical ubiquitin-dependent proteolysis, ubiquitination may interfere with numerous other aspects of the regulation and functions of proteins, such as the intracellular transport of proteins (e.g., Buttner et al., 2001; Schmidt et al., 2021; Lopez-Castejon, 2019; Kwon and Ciechanover, 2017), including of CHTs *in vitro* (Hartnett et al., 2014; Cuddy et al., 2012; Foot et al., 2017).

Second, we investigated the role of immune modulators - cytokines, including two chemotactic members of the cytokine family, the chemokines - in CHT ubiquitination, hypothesizing that, in STs, upregulated levels of immune modulators are associated with elevated CHT ubiquitination levels. To generate mechanistic evidence supporting this relationship, we further hypothesized that increases in cytokine levels evoked by lipopolysaccharide (LPS)-induced activation of the innate immune system cause further CHT ubiquitination. These hypotheses were derived in part from evidence indicating bidirectional interactions between cholinergic activity and anti-inflammatory mechanisms in the brain and periphery (e.g., Hanisch et al., 1993; Leite et al., 2016; Shytle et al., 2004; Rosas-Ballina et al., 2011; Silverman et al., 2015), and between the activity of immune modulators and ubiquitination of immune response-mediating proteins (e.g., Lopez-Castejon, 2019; Lisak et al., 2011; Perga et al., 2021).

CHT ubiquitination and levels of 10 cytokines were determined in two brain regions, the frontal cortex and striatum. The focus on these brain regions reflected the collective evidence indicating that the opponent cognitive-motivational biases of STs and GTs are mediated via variations in dopaminergic and cholinergic signaling in cortical and striatal circuitry (e.g., Campus et al., 2019; Flagel et al., 2011; Pitchers et al., 2017a; Fraser and Janak, 2017). Furthermore, we measured cytokine levels in the spleen, as this organ serves as a primary peripheral source of inflammation-induced increases in immune signaling, including cytokine secretion (del Rey et al., 2009; Murray et al., 2017; Lewis et al., 2019).

The results indicate elevated levels of ubiquitinated CHTs and of several cytokines in the brain, but not spleen, of STs, and that stimulation of the innate immune system further elevates CHT ubiquitination levels in parallel with increases in cytokine production. Together our findings signify a role of neuro-immune modulators in the regulation of the choline transporter, thereby suggesting a role of these modulators in the addiction vulnerability trait indexed by sign-tracking.

## Materials and Methods

### Subjects

Adult Sprague-Dawley rats (N=197; 113 females) between 2 and 3 months of age were purchased from Envigo (Haslett, MI) and Taconic (Cambridge City, IN). Rats weighed 250-275 g upon arrival and were individually housed in Plexiglas cages on a 12 h light/12 h dark cycle (lights on at 7:00 AM), with regulated temperature and humidity (23°C, 45%). After arrival, rats were given one week to acclimate to the colony room before experiments began. Food (Laboratory Rodent Diet 5001; LabDiet) and water were available *ad libitum*. Behavioral testing and experiments were conducted during the light phase (7:00 AM – 7:00 PM). All procedures were approved by the University of Michigan Institutional Animal Care and Use Committee and were conducted in laboratories accredited by the Association for Assessment and Accreditation of Laboratory Animal Care.

Following phenotype assessment, brain tissues and spleen were harvested to determine levels of brain CHT ubiquitination, brain and spleen cytokines and chemokines, and the effects of LPS on these measures. All experiments were designed to follow identical timelines, including animal housing periods, PCA testing periods, and the time between PCA testing and tissue harvesting (detailed below).

### Behavioral phenotyping

#### Apparatus

Prior to the onset of PCA testing, rats were handled daily for 3 days and given ~7 banana-flavored sucrose pellets (45 mg; BioServ). Rats were tested in conditioning chambers (20.5 x 24.1 cm floor area, 20.2 cm high; MED Associates Inc.). Each chamber contained a food magazine port located 2.5 cm above the floor in the center of the intelligence panel, a red house light located on the wall opposite the food magazine port (on throughout training sessions), and a retractable lever (Med Associates) located 2.5 cm to the left or right of the food receptacle and 6 cm above the floor. This retractable lever was illuminated when extended with a white LED light placed inside the lever house. For a lever press to be recorded, a force of ~15 g was needed. The pellet dispenser (Med Associates) delivered one 45 mg banana-flavored sucrose pellet (Bio-Serv) into the food magazine port at a time. A head entry was recorded each time a rat broke the infrared photobeam located inside the food magazine port. Each conditioning chamber was placed in a sound-reducing enclosure and a ventilating fan served as background noise. Data collection was controlled by Med-PC IV Behavioral Control Software Suite.

#### PCA testing and primary measures

One day prior to the onset of PCA testing, rats were placed into the test chamber with the red house light illuminated and the lever retracted. 25 food pellets were delivered in accordance with a variable schedule (VI-30; 0-60 s) to foster the reliable retrieval of pellets from the food magazine port. During PCA testing, trials began by illuminating and extending the lever (CS) for 8 s. Immediately following the retraction of the lever, a single banana pellet was delivered into the magazine port (US). A variable intertrial interval (ITI; 90±60 s) started immediately after the retraction of the lever. An individual PCA training session consisted of 25 trials and lasted 35-40 min. The following 5 measures were extracted from individual trials and collapsed over individual PCA sessions: (1) number of lever deflections (contacts); (2) latency to first lever deflection; (3) number of head entries into the food magazine port (referred to as food cup entries) during the presentation of the CS; (4) latency to the first magazine port entry after CS presentation; and (5) number of magazine port entries during the ITI. Data from animals included in the final analysis reflected sessions during which rats consumed all food pellets.

#### Phenotype classification

The propensity for rats to approach the lever CS versus the food magazine port during the CS period was expressed by a PCA index score. Briefly, the PCA index score consisted of averaging three measures of the conditioned approach on the 4th and 5th day of testing (e.g., Pitchers et al., 2017b; Pitchers et al., 2017a; Yager et al., 2015; Meyer et al., 2012; Robinson and Flagel, 2009): (1) the probability of contacting either the lever CS or food magazine port during the CS period (P(lever) – P(food port)); (2) the response bias for contacting the lever CS or the food magazine port during the CS period: (# lever CS contacts - # food magazine port contacts)/(# lever CS contacts + # food magazine port contacts); and (3) the latency to contact the lever CS or the food magazine port during the CS period: (food magazine port contact latency - lever CS contact latency)/8. Averaging these three measures yields PCA index scores ranging from −1.0 to +1.0, where +1.0 indicates an animal made a sign-tracking CR on every trial and −1.0 a goal-tracking CR on every trial. Further, rats with an averaged PCA index score ranging from −1.0 to −0.5 were classified as GTs, and rats with a PCA index score between +0.5 and +1.0 as STs.

### CHT ubiquitination: TUBEs pull-down assay and immunoblotting

For initial characterization of polyubiquitinated CHTs, tissues were harvested, and frontal cortex and striatum were immediately dissected (tissues from both hemispheres were pooled). Isolated tissues were homogenized in ice-cold 0.32 M sucrose and centrifuged at 1000 x g for 4 min at 4° C to remove cellular debris. The supernatant was centrifuged at 12,500 x g for 15 min to yield a synaptosomal pellet. The pellet was lysed in 0.05 M Tris-HCl buffer containing 1 mM EDTA, 1% Nonidet P-40, 10% glycerol and a protease inhibitor cocktail. Lysates were obtained following centrifugation at 14000 x g for 10 min at 40° C. Protein concentrations in the lysates were determined by using a modified Lowry Protein Assay Kit (Pierce). Pellets were prepared at the University of Michigan and shipped frozen to Temple University where experimenters remained blinded to the phenotypic identity of the samples. We employed the Tandem Ubiquitin Binding Entities (TUBEs) technology to isolate poly-ubiquitinated CHTs (Hjerpe et al., 2009; Yoshida et al., 2015). To confirm isolation, lysates each containing 200 μg of protein were incubated with 20 μL of a slurry containing either Agarose anti-ubiquitin TUBEs (UM402; Life Sensors) or control agarose beads (UM400; Life Sensors) that were pre-equilibrated in working buffer (0.05M Tris-HCl containing 0.1% Tween 20) on a shaker overnight at 40° C. The samples were spun twice at 5000 x g to remove the supernatant and the residue (pull-down bound fraction) containing the beads was processed for immunoblotting. Pull-down samples were resuspended in Laemmli buffer and proteins were separated on 4–15% Tris HCl polyacrylamide gels using electrophoresis and transferred to the PVDF membranes. Immunodetection of ubiquitinated proteins was accomplished by incubating the membrane overnight with 1:1000 diluted monoclonal ubiquitin antibody P4D1 (sc-8017; Santa Cruz Biotechnology) and exposing the membranes to peroxidase-conjugated anti-rabbit secondary antibody and ECL Advance chemiluminescent substrate (GE Healthcare). Poly-ubiquitinated CHTs were detected by stripping the membranes with Restore Plus buffer (Pierce) and incubating them with 1:2000 diluted rabbit anti-CHT polyclonal antibody (ABN458; EMD Millipore). The resulting chemiluminescent signal was acquired with a Chemidoc Touch Imaging System (Bio-Rad). After confirming the efficacy of Agarose TUBEs to isolate and detect poly-ubiquitinated CHTs, all subsequent subcellular fractionation studies used lysates containing 100 μg of proteins to isolate and compare the densities of poly-ubiquitinated CHTs in ST and GT samples. Because it was not possible to normalize densities with a control protein in pull-down samples, we ran separate gels with lysates (25 μg protein) to determine the densities of β-actin protein. Densitometric analysis of poly-ubiquitinated CHT-immunoreactive bands was performed by calculating the integrated pixel densities using NIH ImageJ software and the data were normalized to β-actin densities.

To determine whether the distribution of poly-ubiquitinated CHTs differed between different cellular domains, we employed a subcellular fractionation strategy to isolate the plasma membrane-enriched (LP1) and vesicular membrane-enriched (LP2) fractions as described previously (Ferguson et al., 2003; Apparsundaram et al., 2005; Parikh et al., 2013; Koshy Cherian et al., 2017). Briefly, the synaptosomal pellet was lysed in 5 mm HEPES-NaOH under ice-cold conditions. LP1 fraction was collected by spinning the lysate at 15,000 × g for 20 min. The resultant supernatant was centrifuged at 200,000 × g for 30 min to obtain the LP2 fraction using a TL100 ultracentrifuge (Beckman Coulter). Both LP1 and LP2 pellets were resuspended in Tris-HCl buffer containing 1mM EDTA, 1% Nonidet P-40, 10% glycerol and protease inhibitor cocktail. Following protein estimations, samples from both fractions containing 100 μg were subjected to TUBEs pull-down procedure followed by immunoblotting as described above.

As described in Results, ubiquitinated CHTs in the LP2 fraction largely accounted for phenotypic differences in poly-ubiquitinated CHTs. Based on these initial findings, subsequent analyses of the effects of LPS challenge on levels of ubiquitinated CHTs were conducted in total synaptosomes. Methods for the administration of LPS and subsequent tissue harvesting were identical to these described in the experiments measuring cytokines and chemokines and are detailed below.

### Administration of LPS and LPS-induced sickness behavior

In addition to the determination of brain CHT ubiquitination levels and concentrations of cytokines and chemokines (below) in STs and GTs, the effects of activation of the innate immune system by the bacterial endotoxin lipopolysaccharide (LPS) were assessed. LPS, from Escherichia coli (serotype O111:B4), was obtained from Millipore Sigma (St. Louis, MO) and suspended in 0.9% saline (Teknova 0.9% sterile saline solution; Fisher Scientific; Hollister, CA) vortexed, allocated to 1.0 mg/mL, and stored in 1.5 mL Eppendorf tubes at −20°C until used. Preparation and injections of LPS were conducted in a chemical fume hood and using procedures approved by the University of Michigan Environment, Health & Safety Department.

STs and GTs were randomly assigned to the administration of LPS (1.0 or 5.0 mg/kg in 0.2 mL saline; i.p) or 0.9% saline. Sickness behaviors, including piloerection, ptosis, lethargy, and huddling were monitored each hour for six hours following the treatment injection. A lethal dose of sodium pentobarbital was administered (1.5 g/kg; i.p.) 6 hrs after injections, followed by transcardiac perfusion with 0.9% saline. The right frontal cortex and right striatum were dissected out and synaptosomal pellets were prepared for determination of CHT ubiquitination levels as described above. Furthermore, spleen, left frontal cortex, and left striatum were harvested, flash-frozen in 2-methyl butane, and stored at −80°C for subsequent determination of cytokine and chemokine levels.

#### Sickness behavior

Four symptoms of sickness behavior (piloerection, ptosis, lethargy, and huddling; e.g., Kelley et al., 2003) were monitored and recorded each hour for six hours following the administration of LPS (1.0 mg/kg) and vehicle in 8 rats per treatment condition (4 per phenotype). The researchers observing sickness behavior were blinded to the phenotype. At each time point, the presence of absence of these four symptoms was recorded. Piloerection was marked when the animals’ body hair stood up, eyelid drooping was scored as ptosis, lethargy was scored if a rat did not approach the front of the home cage when approached by the researcher, and hunched or curled body posture was marked as huddling.

### Cytokine and chemokine magnetic bead panel

Multiplex immunobead assay technology (Milliplex MAP Rat Cytokine/Chemokine Magnetic Bead Panel, Millipore Corp., St. Louis, MO; Magpix Analytical Test Instrument, Luminex Corp., Austin, TX) was used to determine brain and spleen cytokine levels (e.g., Hulse et al., 2004; Erickson and Banks, 2011; Fox et al., 2005). The previously flash-frozen tissues were homogenized in assay buffer (Milliplex Rat Cytokine Kit) and protein concentrations were determined using the Bradford Assay and an Epoch BioTek Microplate Spectrophotometer (Gen 5 2.01; BioTek Instruments Inc; Winooski, VT). 25 μg of protein from tissue homogenates were combined with specific cytokine antibody-immobilized beads, mixed with bead dilutant, and incubated at room temperature for 2 hrs on a plate shaker. The well contents were washed, detection antibodies μwere added, and the plate was incubated at room temperature for 1 hr on the plate shaker. Without aspirating the contents of each well, streptavidin-phycoerythrin was added and the plate was incubated at room temperature for 30 min on the plate shaker. Contents of the well were then removed, washed, and drive fluid was added to resuspend the beads. The MAGPIX plate reader (xPONENT software; Luminex Corporation; Austin, TX) was used to measure quantities of fluorescent-coded magnetic beads and determine concentrations of selected cytokines: interleukin 1-alpha (IL-1α), interleukin 1-beta (IL-1β), interleukin-6 (IL-6), interleukin-4 (IL-4), interleukin-2 (IL-2), interleukin-18 (IL-18), interferon-gamma (IFN-γ), C-C motif chemokine ligand 2 (CCL2) and C-X-C motif chemokine ligand 10 (CXCL10). A calibration curve was generated to determine each sample’s cytokine concentration. Readings that were below detectable levels of analyte were recorded as the lowest possible measurement value. Furthermore, approximately 10% of samples did not contain a sufficient concentration of proteins (<1.0 μg/μL) to determine cytokine levels by the MAGPIX plate reader. These samples were excluded from the analysis, resulting in a variable number of data points for cytokine levels per brain region or spleen and phenotype.

### Experimental design and statistical analysis

Differences between CHT ubiquitination levels of STs and GTs were analyzed using two-sided t-tests. For comparisons across subcellular fractions, obtained from the same total synaptosomal preparation, alpha was set at 0.05/2. Basal cytokine levels in STs and GTs were compared using non-parametric Mann-Whitney tests because for several analytes, all, or nearly all, measures from GTs were at the assay’s detection threshold and thus data were not normally distributed. The effects of LPS (vehicle and two doses) on CHT ubiquitination levels were analyzed using 2-way ANOVAs on the effects of treatment and phenotype, followed by one-way ANOVAs (where applicable) and pairwise comparisons (uncorrected Fisher’s LSD test) as permitted by ANOVA results. For parametrically analyzed data, graphs depict individual values, means and 95% Confidence Intervals (CI). Statistical analyses were performed using SPSS for Windows (version 17.0; SPSS) and GraphPad Prism (version 9.4.1). Exact *P* values were reported (Greenwald et al., 1996; Sarter and Fritschy, 2008; Michel et al., 2020). For major results derived from parametric tests, effect sizes (Cohen’s *d*) were indicated (Cohen, 1988).

## Results

### Screening of STs and GTs

PCA testing was conducted in a total of N=197 rats (113 females). PCA scores from the 4^th^ and 5^th^ sessions were averaged for individual rats, and this average score determined phenotype assignment (GT, ST, INT; PCA scores of ±0.5 served as cut-off points for the classification of STs and GTs).

Individual PCA scores obtained from the 1st, 4^th^ and 5^th^ PCA session are shown in Fig. 1. During the first session, rats did not exhibit a preference for approaching and operating the lever (i.e., the CS) versus entering the magazine, as reflected by a mean PCA score near zero. Across subsequent testing sessions, an increasing number of rats developed a preference for the CS or the magazine, yielding 87 STs (56 females). 43 GTs (20 females) and 67 INTs (37 females). A Fisher’s exact test was used to examine the sex-based distribution of STs and GTs (*P*=0.06). This insignificant trend reflected that females were more likely to be classified as STs. Although a similar pattern was previously observed (Kucinski et al., 2022), it has not been consistently found (Pitchers et al., 2015), reflecting considerable variation in the sex-dependent distribution of these phenotypes across cohorts obtained from different vendors as well as from different breeding barriers from the same vendors, and associated genetic heterogeneity (Fitzpatrick et al., 2013; Gileta et al., 2022). Note that in Fig. 1, individual PCA scores from days 4 and 5 were graphically assigned to phenotypes based on the final classification (as opposed to the classification that would be based on the scores from just one session).

**Figure 1.**
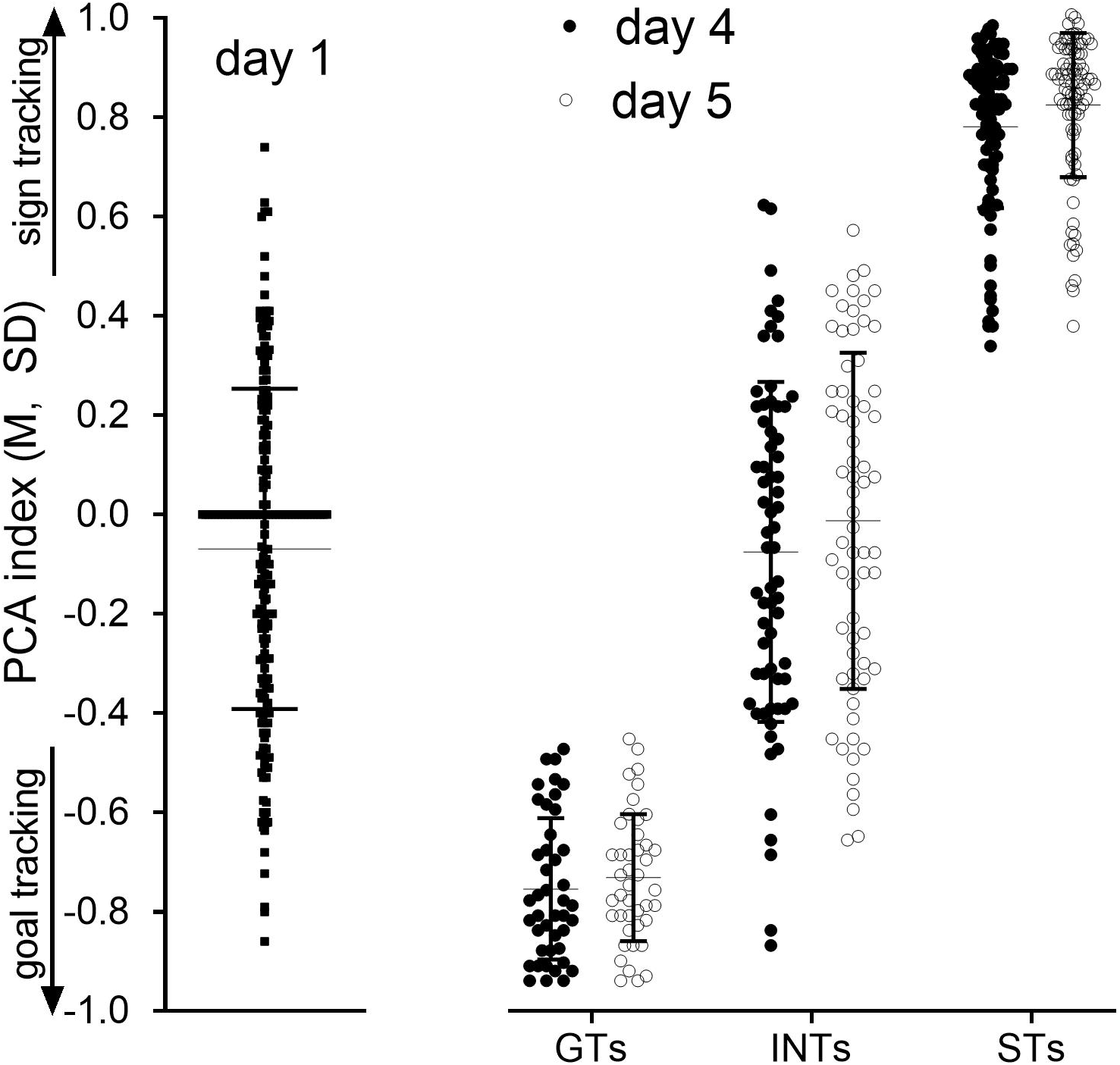
Behavioral screening of STs and GTs using the Pavlovian Conditioned Approach (PCA) test (shown are individual data points, means and SDs; note that the thicker bar at the day 1 PCA index of 0.0 represents a cluster of individual data points). The behavior of a total of N=197 rats (113 females) was assessed across 5 consecutive PCA sessions (over 5 days). PCA scores reflected whether rats preferably approached and contacted the lever (Pavlovian food cue; positive PCA scores; STs) or approached the magazine (negative PCA scores; GTs). Scores obtained from the first session (left) indicated that most rats did not exhibit a bias toward the lever (CS) or the magazine (note that the thick horizontal bar depicts multiple data points with PCA scores at zero). By days 4 and 5 (right), nearly a third of the rats had developed a preference either for approaching the CS or emerged as GTs (the graph shows PCA scores from test days 4 and 5, assigned to the three phenotypes based on the final classification of phenotype using the average of individual PCA scores obtained from sessions 4 and 5; cut off scores: ±0.5). Rats with intermediate PCA scores (INTs) were not further investigated.

### Elevated levels of poly-ubiquitylated CHT in STs

The relatively poor attentional performance of STs, and the associated propensity for approaching and contacting a Pavlovian cocaine cue, were previously attributed to attenuated levels of cortical cholinergic signaling when compared with GTs (Paolone et al., 2013; Pitchers et al., 2017a; Pitchers et al., 2017b; Pitchers et al., 2018). Furthermore, a deficient translocation of CHTs, from intracellular domains into synaptosomal plasma membrane, was found to underly the dampened capacity of the cholinergic system in STs (Koshy Cherian et al., 2017). Here, we first explored a posttranslational CHT modification – poly-ubiquitination – as a potential mechanism contributing to the deficient outward trafficking of CHTs in STs. Ubiquitination proteolytically cleaves synaptic proteins or interferes with trafficking of such proteins, including the CHT (Buttner et al., 2001; Hartnett et al., 2014; Meyer et al., 1986; Yamada et al., 2012).

We first measured CHT ubiquitination in a total synaptosomal preparation extracted from frontal cortex of STs and GTs. Figure 2a documents the efficiency of the TUBEs method and Figure 2b exemplifies the lower levels of ubiquitinated CHTs in frontal cortex of an ST compared to a GT. Levels of ubiquitinated CHTs in total synaptosomal lysates from cortex of STs were significantly higher in STs (t(8)=2.66, *P*=0.028; Cohen’s *d*: 1.68; Fig. 2c). Subsequently, and based on tissues obtained from a separate cohort of STs and GTs, we employed the TUBEs method to isolate poly-ubiquitinated CHTs separately from surface membrane- and vesicular membrane-enriched synaptosomal fractions. Ubiquitination levels of CHTs residing in the synaptosomal plasma membrane were extremely low in both phenotypes, approaching the detection limit of our assay (t(9)=0.17, *P*=0.87; Fig 2d, left). In contrast, concentrations of ubiquitinated CHTs located in intracellular domains were greatly elevated in STs over GTs, with no overlap between the individual data from the two groups (t(9)=4.94, *P*=0.008; Cohen’s *d*: 2.68; Fig. 2d, right). Ubiquitination of intracellular CHTs in STs may be causally related to the attenuated outward trafficking in response to increases in neuronal activity (Koshy Cherian et al., 2017).

**Figure 2.**
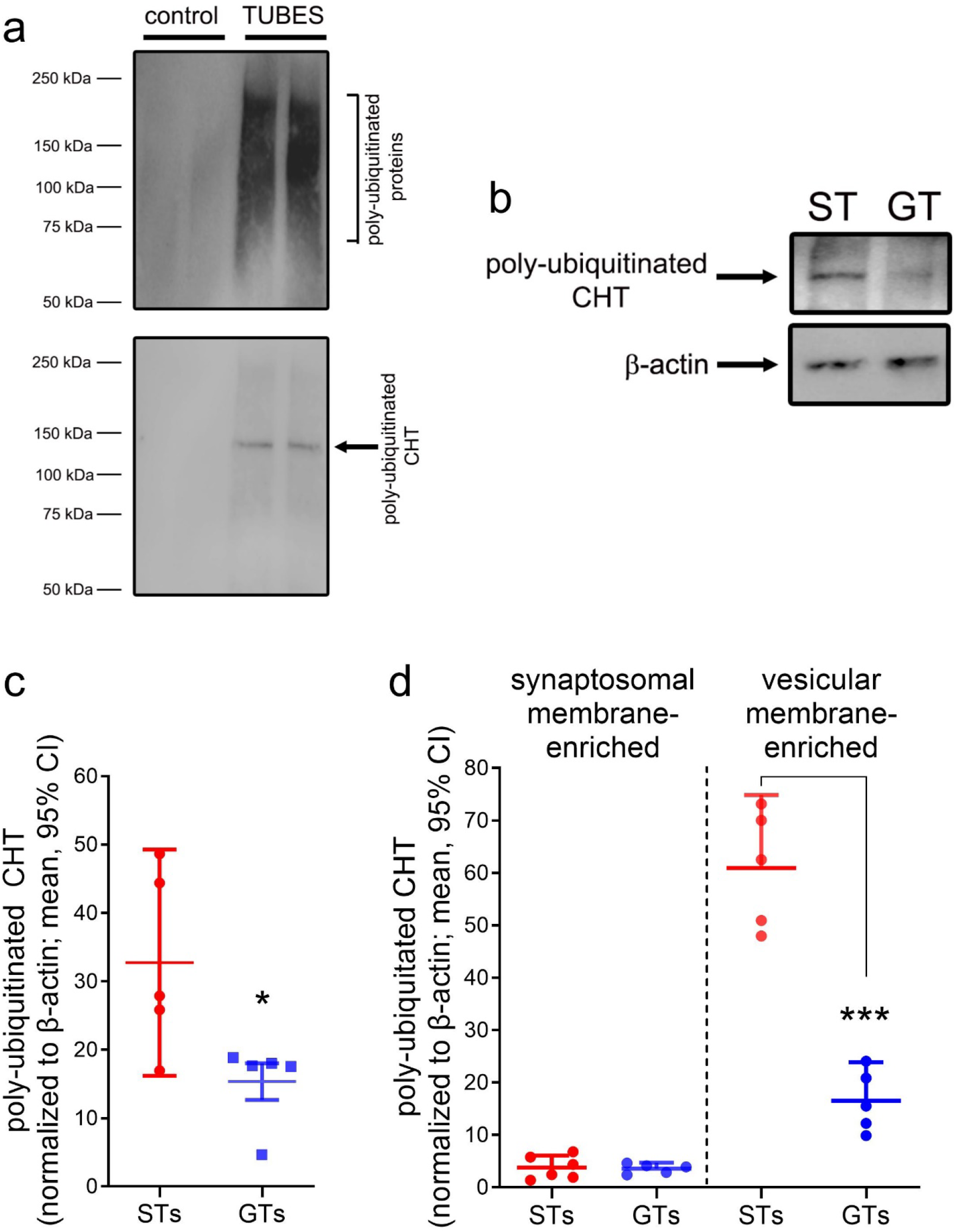
Levels of ubiquitinated CHTs in the cortex of STs and GTs. <u>a</u> Demonstration of the efficacy of the TUBEs technology to isolate poly-ubiquitinated CHTs. Upper panel: Immunoblotting of the lysates from 2 STs incubated with agarose TUBEs reveal a strong enrichment of poly-ubiquitinated proteins (range: 50-250 kDa). Lower panel: Re-probing membranes with the anti-CHT antibody detected a poly-ubiquitinated CHT band in a higher molecular weight range (100-150 kDa) than the unconjugated CHT that is typically observed at ~55 kDa (Koshy Cherian et al., 2017). A chemiluminescence signal was not detected from synaptosomal lysates incubated with control agarose beads. <u>b</u> Immunoblot depicting poly-ubiquitinated CHT proteins isolated from frontal cortex synaptosomal lysate of a representative ST and GT rat. <u>c</u> Levels of ubiquitinated CHTs in total synaptosomal lysates from cortex of STs were significantly higher than in GTs (5 rats per phenotype; unless noted otherwise, this and subsequent data figures show individual data points, means and 95% confidence intervals (CI); symbols used to indicate results from *post hoc* multiple comparisons or two-group comparisons: *,**,***” *P*<0.05, 0.01, 0.001). <u>d</u>: A subsequent determination of CHT ubiquitination levels in subcellular fractions (isolating synaptosomal membrane- and vesicular membrane-enriched fractions) indicated the near-absence of ubiquitinated CHTs in synaptosomal membrane-enriched fractions prepared from either phenotype (prepared form a separate cohort of 5 rats per phenotype). In contrast, levels of ubiquitinated CHTs from intracellular, vesicular membrane-enriched fractions were robustly higher in STs than in GTs (note that there is no overlap in the individual data points between the two groups).

### Elevated brain, but not spleen, cytokine levels in STs

Cholinergic activity influences neuroinflammatory activity in the brain and periphery (e.g., Lehner et al., 2019; Martelli et al., 2014; Rosas-Ballina et al., 2011). The concept of cholinergic-anti-inflammatory pathways (Martelli et al., 2014) predicts that elevated levels of neuroinflammation result from a dampened or dysfunctional cholinergic system (e.g., Leite et al., 2016; Rosas-Ballina and Tracey, 2009; Field et al., 2012). Moreover, bidirectional interactions between elevated levels of ubiquitination and neuroinflammatory mechanisms have been demonstrated (Lopez-Castejon, 2019; Schmidt et al., 2021), further suggesting the possibility that, in STs, brain and peripheral cytokine and chemokine levels are upregulated and contribute to ubiquitination of CHTs.

We measured levels of a total of 10 immunomodulators in two brain regions (frontal cortex, striatum) and spleen. Four of these 10 analytes have traditionally been considered to exert primarily proinflammatory effects (IL-1α, IL1-β, IL-2, IL-6, IFN-γ; red background in Figs. 3 and 5), 3 cytokines have been viewed as predominantly anti-inflammatory (IL-4, IL-10 IL-18; green background), and the remaining two analytes were chemokines (CCL2, CXCL 10; e.g., Turner et al., 2014).

**Figure 3.**
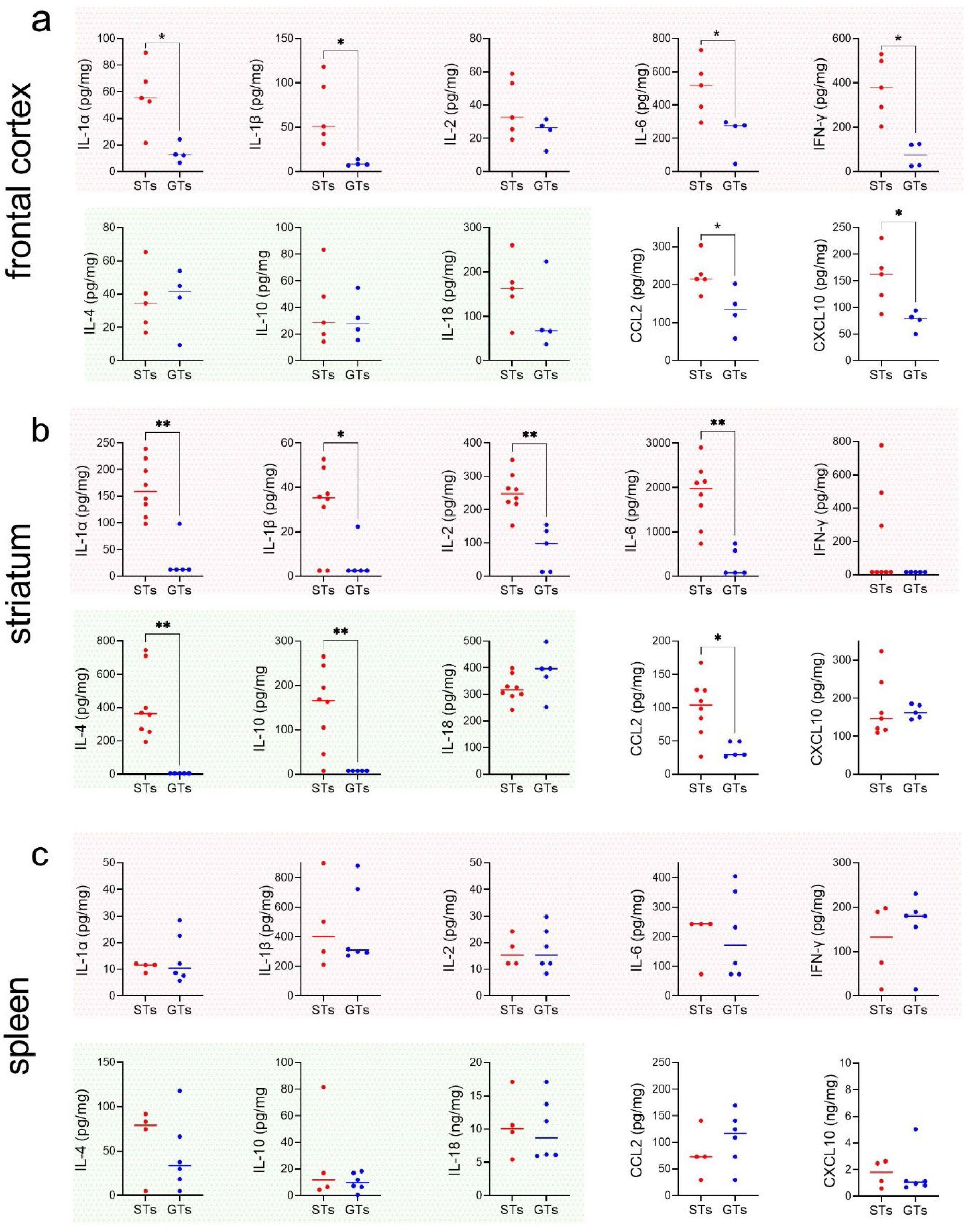
Elevated levels of cytokines and chemokines in the brain, but not spleen, of STs when compared with GTs (shown are individual values and medians; frontal cortex: n=5 STs, 3 females; 4 GTs, 1 female; striatum: 8 STs, 6 females. 5 GTs, 1 female; spleen: 4 STs, 3 females, 6 GTs, 2 females; variable numbers of data points per phenotype, brain region and spleen resulted from tissue handling issues and assay failures). Levels of traditionally considered pro-inflammatory cytokines (plots with red background), anti-inflammatory cytokines (green background), and two chemokines (CCL2 and CXCL2) were measured from frontal cortical tissue (<u>a</u>), striatal tissue (<u>b</u>), and spleen (<u>c</u>). Indications of significant differences between the phenotypes are based on Mann-Whitney U tests (*,**: *P*<0.05, 0.01). Note that significant phenotype effects based on values obtained from striatal tissues in GTs (<u>b</u>) were at or below detection threshold. This observation, together with the contrasting absence of any significant differences between cytokine and chemokine levels in spleen, supports the robustness of the finding that the majority of cytokines and chemokines measured in brain tissues were upregulated in STs when compared with GTs.

**Figure 4.**
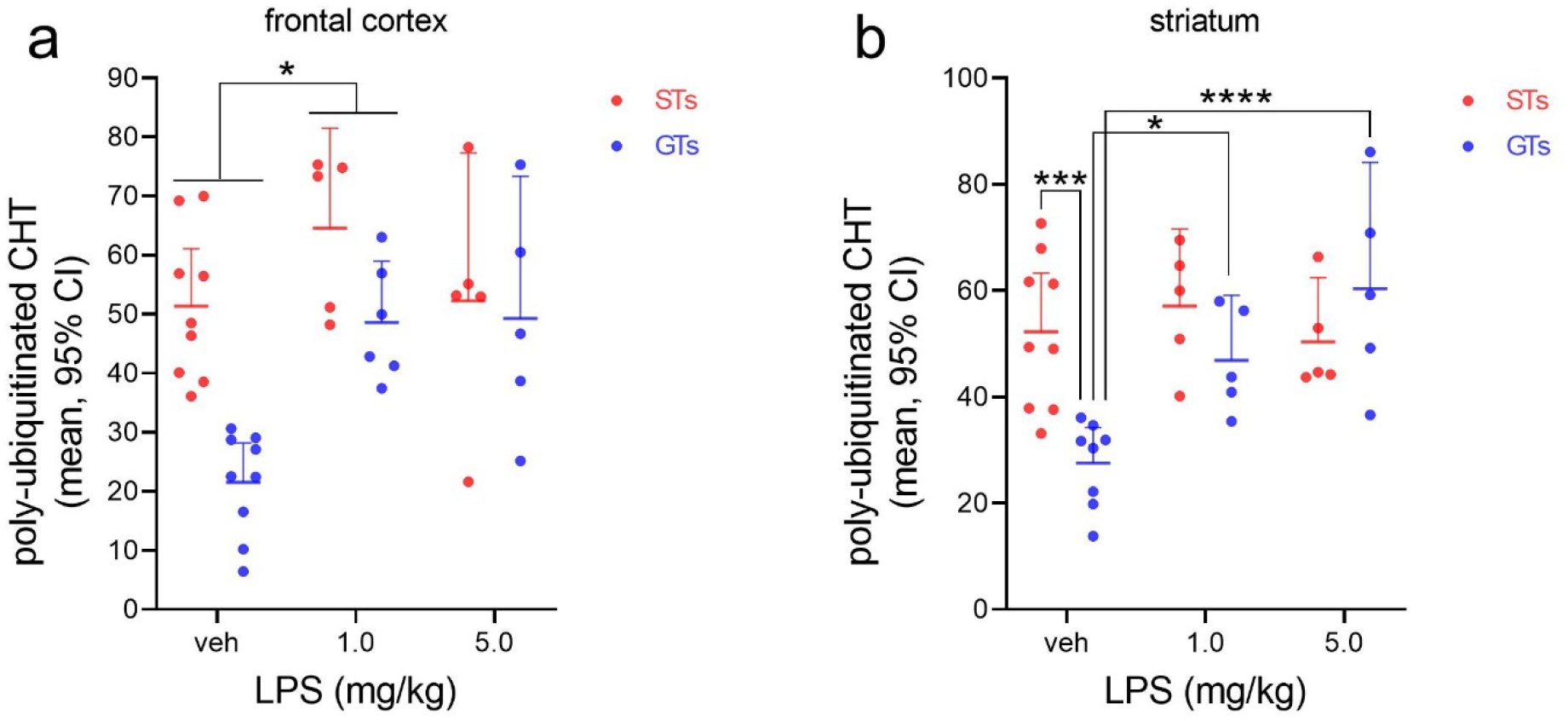
Administration of LPS increased ubiquitination levels of CHTs prepared from frontal cortical (<u>a</u>) and striatal (<u>b</u>) tissues (19 STs, 9 treated with vehicle (2 females), 5 with 1.0 mg/kg LPS (2 females) and 5 with 5.0 mg/kg LPS (3 females); 20 GTs, 9 treated with vehicle (3 females), 6 with 1.0 mg/kg LPS (1 female) and 5 with 5.0 mg/kg LPS (2 females)). Analysis of effects of LPS on cortical ubiquitination levels indicated main effects of phenotype and treatment, while the interaction between the effects of phenotype and treatment failed to reach significance (*P*=0.055). The main effects reflected higher levels of ubiquitinated CHTs in STs (see Results) and increases in levels after LPS (*post hoc* multiple comparisons indicated a significant increase following the lower dose of LPS when compared with controls; <u>a</u>). The effects of LPS and phenotype on striatal CHT ubiquitination interacted significantly, allowing the multiple comparisons across groups and treatments (results indicated in <u>b</u>). In rats treated with vehicle, elevated levels of ubiquitinated CHTs were again found in STs when compared with GTs. LPS increased levels of CHT ubiquitination only in GTs, abolishing the phenotype effects (and mirroring effects of LPS on cortical levels). In neither region did the higher dose of LPS produce a greater effect than the 1.0 mg/kg dose. Together, these data suggest that baseline CHT ubiquitination levels are at a maximum in STs that cannot be further activated by this endotoxin. The efficacy of LPS to ubiquitinate CHTs is consistent with the overall hypothesis that CHT regulation interacts with neuroinflammatory signaling.

**Figure 5.**
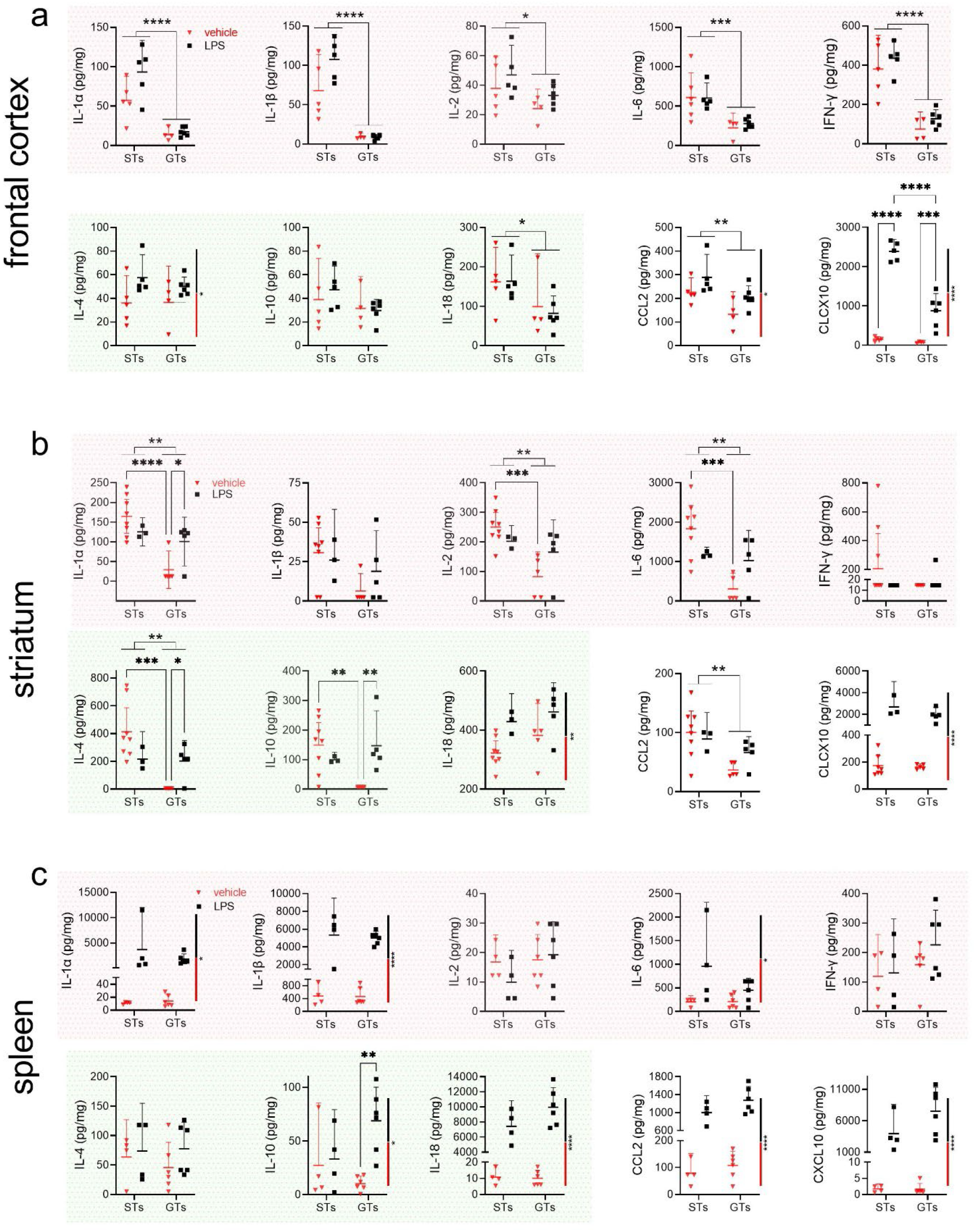
Effects of LPS on brain and spleen cytokine and chemokine levels (cortex: 10 STs, 5 treated with vehicle (3 females) and 5 treated with 1.0 mg/kg LPS (2 females); 10 GTs, 4 treated with vehicle (1 female) and 6 treated with LPS (1 female); striatum: 11 STs, 8 treated with vehicle (6 females) and 3 treated with LPS (3 females); 10 GTs, 5 each treated with vehicle (1 female) and 5 treated with LPS (1 female); spleen: 8 STs, 4 treated with vehicle (3 females) and 4 treated with LPS (3 females); 12 GTs, 6 treated with vehicle (2 females) and 6 treated with LPS (1 female)). Note that while this figure resembles the design of Figure 3, the abscissa of this figure depicts phenotype, to graphically emphasize the effects of LPS per phenotype. Main effects of phenotype and the results of multiple comparisons (Fisher’s LSD), if permitted by significant interactions between the effects of phenotype and LPS treatment, are indicated by data column-connecting lines and stars (*,**,***” *P*<0.05, 0.01, 0.001). Main effects of treatment are indicated by vertical, black-red bars at the right side of individual graphs (see most graphs of data from spleen). LPS increased cytokine and chemokine levels of brain chemokines and IL-4 and IL-18 in cortex (<u>a</u>) and striatum (<u>b</u>), respectively. Phenotype-selective effects on brain cytokine were restricted to increases in GTs (3 cytokines in striatum) and generally not observed in STs. Most analytes measured from spleen (<u>c</u>) showed large LPS-induced increases in both phenotypes exemplifying, as expected, the relatively superior efficacy of the endotoxin on spleen cytokine levels when compared with brain. Only brain CXCL10 concentrations showed comparably large increases following LPS.

In frontal cortex (Fig. 3a) and striatum (Fig. 3b), concentrations of 6 and 7 analytes, respectively, were significantly higher in STs than GTs, including 4 out of 5 pro-inflammatory cytokines in each region. Among traditionally classified anti-inflammatory cytokines, IL4 and IL-10 levels were significantly higher in the striatum of STs. Levels of both chemokines were significantly higher in the cortex of STs compared with GTs while, in the striatum, this was the case only for CCL2. In spleen (Fig. 3c), neither cytokine nor chemokine levels differed between the phenotypes.

As detailed in Methods, cytokine measures at or below the detection threshold of our assays were assigned the lowest measurable concentration value. In the striatum, among the 7 analytes that yielded significant differences between the phenotypes, nearly all (5 out of 7) measures in GTs were at or below the detection threshold (IL-1α, IL1-β, IL6, IL4, IL-10). In contrast, in STs, levels of these analytes reached ~40-2000 pg/mg, further indicating the significance of the phenotype effect on brain cytokine levels. Given that PCA scores of STs and GTs were densely clustered within the range of scores underlying phenotype classification, and that cytokine levels, particularly in GTs, often compressed toward detection limits, the two measures were not significantly correlated (Pearson’s r; STs: all r<0.75, all *P*>0.13; GTs: all r<0.68, all *P*>0.31).

### LPS increased CHT ubiquitination

The evidence described above indicated that cortical CHTs of STs are highly ubiquitinated and that levels of many cytokines are upregulated in the brain, but not spleen, of STs when compared with GTs. Given that ubiquitination of CHTs in STs was entirely restricted to those located in intracellular domains, it appears likely that this modification contributes to the disruption of neuronal activity-associated CHT outward trafficking and the associated attenuation of cholinergic activity characteristic of this phenotype (Koshy Cherian et al., 2017; Pitchers et al., 2017b; Pitchers et al., 2017a). *Vice versa*, the presence of upregulated neuro-immune modulators in STs may contribute to the ubiquitination of CHTs, giving rise to reciprocal interactions between disrupted cholinergic signaling, elevated neuroinflammatory signaling and CHT dysregulation (e.g., Lopez-Castejon, 2019; Lisak et al., 2011). To determine a causal mechanism integral to these bidirectional relationships between CHT status and cytokine levels, we tested the hypothesis that administration of the bacterial endotoxin LPS, which has been extensively documented to produce a pleiotropic set of sickness responses, including elevated cytokine release (e.g., Datta and Opp, 2008; Erickson and Banks, 2011), increases CHT ubiquitination. Given elevated CHT ubiquitination levels at baseline in STs (above), we also expected to see relatively greater effects of LPS on CHT ubiquitination levels in STs. Because the evidence described above located elevated CHT ubiquitination levels in STs solely in the intracellular domain, and the technical challenges associated with measuring CHT ubiquitination in subcellular fractions, we determined ubiquitination levels in total synaptosomal preparations and attributed all CHT ubiquitination to CHTs located in intracellular domains.

Administration of LPS elicited comparable sickness behaviors in STs and GTs (Table 1). Hourly observations (by a blinded research assistant) of the presence of piloerection, lethargy, huddling and ptosis indicated a highly consistent course of the sickness response in a subgroup of STs and GTs. Two of the 4 symptoms were present in all rats beginning 1 hr following the injection, and all 4 symptoms were observed in all rats from hour 3 to hour 6 post-injection. Vehicle-treated rats did not show any of these symptoms. Six hours following the administration of vehicle or LPS (1.0 or 5.0 mg/kg), tissues were harvested. CHT ubiquitination levels in synaptosomes from frontal cortical and striatal tissue of STs and GTs are illustrated in Fig. 4.

**Table 1.**
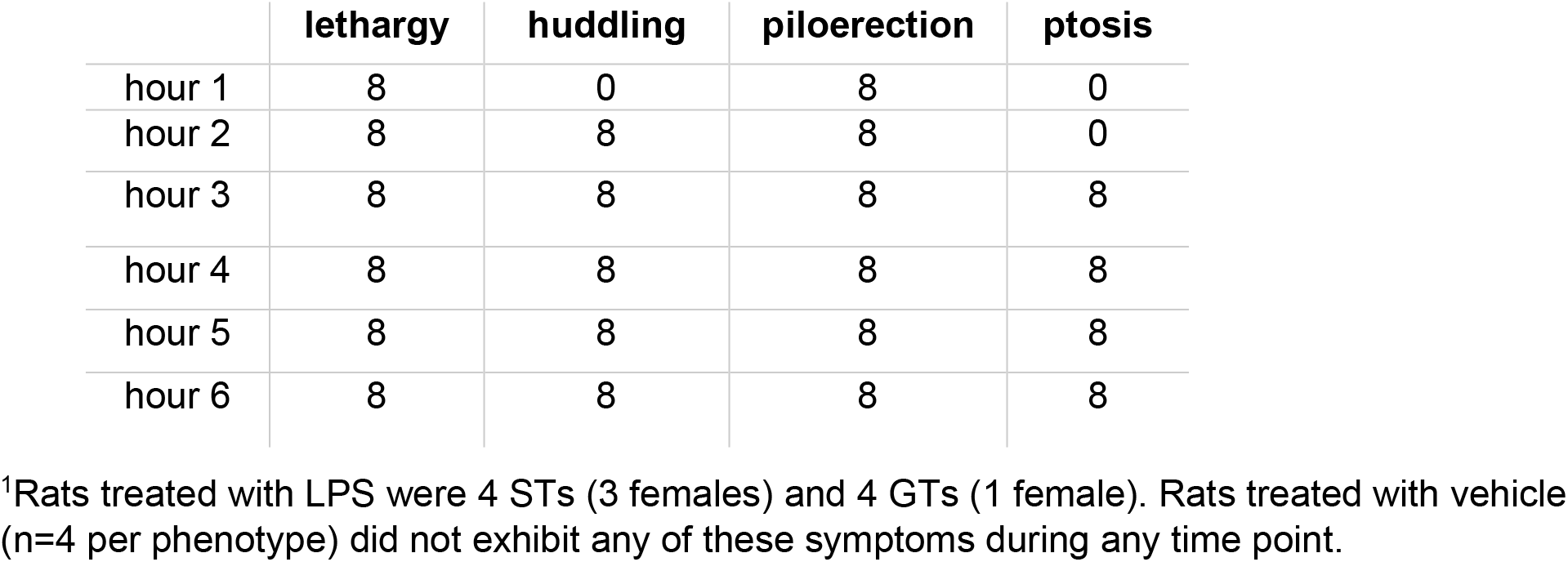
Number of rats exhibiting symptoms of sickness behavior following the administration of LPS (1.0 mg/kg).^1^.

For values from frontal cortex (Fig. 4a), a two-way ANOVA on the effects of phenotype and treatment indicated significant main effects of phenotype (F(1,33)=12.72, *P*=0.0011) and treatment (F(2,33)=8.16, *P*=0.0013), while the interaction between these two factors failed to reach significance (F(2,33)=3.16, *P*=0.056). The main effect of phenotype again reflected that ubiquitinated CHT levels were significantly higher in STs (t(37)=3.41, *P*=0.0016; M, SEM; STs: 55.05±3.53; GTs: 36.55±4.076). Furthermore, a *post hoc* 1-way ANOVA on the effects of LPS (F(2,36)=4.81, *P*=0.041) indicated a significant increase ubiquitinated CHTs after 1.0 mg/kg, but not 5.0 mg/kg, LPS (Fig. 4a). Compared with the effects of 1.0 mg/kg LPS, the higher dose of did not further increase CHT ubiquitination levels (Fig. 4a). Although inspection of the data indicated that LPS increased ubiquitinated CHT levels exclusively in GTs, the absence of a significant interaction term prevented *post hoc* multiple comparisons on the effects of dose on levels in the two phenotypes.

For the values derived from striatal tissues, a significant interaction between the effects of phenotype and treatment (F(2,31)=6.07, *P*=0.006) permitted such multiple comparisons (main effect of treatment: F(2,31)=5.71, *P*=5,71 *P*=0.008; main effect of phenotype: F(2,31)=3.83, *P*=0.059; Fig. 4b). Following vehicle administration, CHT ubiquitination levels again were higher in STs than GTs (Fisher’s LSD: *P*=0.0003; Cohen’s *d*: 2.13), but this difference was abolished after LPS administration (all *P*>0.10), reflecting that both doses of LPS increased ubiquitinated CHT levels in GTs, but not STs (GTs: both *P*<0.001; STs: both *P*>0.49; Fig. 4b). Again, the higher dose of LPS did not further increase effects relative to the lower dose (Fig. 4b).

We also explored potential relationships between PCA scores and CHT ubiquitination levels in rats treated with vehicle. The absence of such correlations in cortical and striatal ubiquitination scores from STs and GTs (Pearson’s r; all r<0.31, all *P*>0.44) may reflect the closely clustered PCA scores within the classification range for either phenotype, or that such classification yields a categorical distinction of traits.

Although the interaction between the effects of LPS and phenotype reach statistical significance for the data from striatum but not from frontal cortex, the evidence from both brain regions together suggests that LPS administration increased CHT ubiquitination primarily in GTs, but not in STs. This finding may reflect that, in STs, levels of CHT ubiquitination at baseline were already at a maximum that cannot be further increased by this pro-inflammatory challenge. The findings that a relatively high dose of LPS (5.0 mg/kg) did not further increase CHT ubiquitination levels relative to the lower dose, and that baseline levels in STs corresponded with the levels seen after this high dose of LPS in GTs, further support this interpretation. The finding that LPS treatment increased levels of CHT ubiquitination may indicate a mechanistic interaction between the activity of neuro-immune modulators and CHT status.

### LPS increased cytokine levels

If cytokine and chemokine levels indeed influence CHT ubiquitination, then the evidence described above may suggest that cytokine levels likewise are already at a “ceiling” in STs, and that LPS-induced increases in cytokine levels therefore may largely be restricted to GTs.

As detailed in Methods and paralleling the experiments on the effects of LPS on CHT ubiquitination levels, tissues were harvested 6 hours following vehicle or LPS treatment. Variations in animal numbers per analyte and brain region and spleen were solely based on technical variables related to tissue handling and various assay failures. Furthermore, and in contrast to Fig. 3, the individual graphs in Fig. 5 place the phenotypes on the abscissa to graphically emphasize the presence of potential LPS effects in each phenotype. Fig. 5 shows the results of multiple comparisons based on 2-way ANOVAs (phenotype x LPS); main effects of LPS are graphically indicated by a vertical black-white bar placed on the right of the individual plots (see, e.g., 7/10 graphs for spleen).

The results of the ANOVAs confirmed the presence of higher levels of most brain cytokines and chemokines in STs relative to GTs (main effects of phenotype; Fig. 5). LPS produced the most widespread and phenotype-independent elevations of analyte levels in spleen (Fig. 5c).

Except for CCL2 concentrations in striatum, LPS most robustly increased brain levels of the two chemokines, irrespective of phenotype (main effects; vertical bars right to individual plots in Fig. 5a and 5b). The concentration of only one additional brain analyte - IL-18 in striatum - was increased by LPS irrespective of phenotype (main effect). In the brain, LPS increased the concentrations of 3 striatal analytes exclusively in GTs (IL-1α, IL-4, IL-10; reflecting significant interactions between effects of phenotype and LPS). Effects restricted to GTs may have paralleled the ceiling effects seen in the analysis of CHT ubiquitination levels (above). In no case did LPS increase the concentration of an analyte solely in STs.

Taken together, the effects of LPS indicated that LPS robustly increased spleen cytokine levels in both phenotypes and, in cortex, LPS produced large increases in the concentrations of the two chemokines. The discussion below will focus on the significance of the result that the concentrations of the chemokines in cortex were higher in STs than in GTs at baseline and also further elevated by LPS.

## Discussion

The present experiments generated five main results (see Fig. 6 for a schematic illustration and summary of these results). 1. Levels of ubiquitinated CHTs, measured in total synaptosomal preparations from cortex, were significantly higher in STs when compared with GTs. 2. Measuring ubiquitinated CHTs in synaptosomal subcellular fractions, CHTs situated in surface plasma membrane were nearly completely devoid of ubiquitination in both phenotypes. In contrast, CHTs detected in vesicular membrane-enriched fractions were significantly more ubiquitinated in STs, with no overlap between individual data points from the two phenotypes (upper left half of Fig. 6). 3. Cytokine levels in spleen did not differ between STs and GTs; in contrast, cortical and striatal concentrations of numerous cytokines and chemokines were significantly higher in STs when compared to GTs (lower left in Fig. 6). 4. Systemic administration of LPS increased levels of ubiquitinated CHTs in GTs, but not STs. Following LPS, CHT ubiquitination levels in GTs were statistically similar to those measured in STs (upper right in Fig. 6). 5. LPS increased levels of nearly all cytokines in spleen, irrespective of phenotype. Two cytokines and two chemokine were increased after LPS (main effects). Of these, the two chemokine were also higher in STs at baseline (lower right in Fig.6). In no case did LPS increase the concentrations of a cytokine or chemokine exclusively in STs.

**Figure 6.**
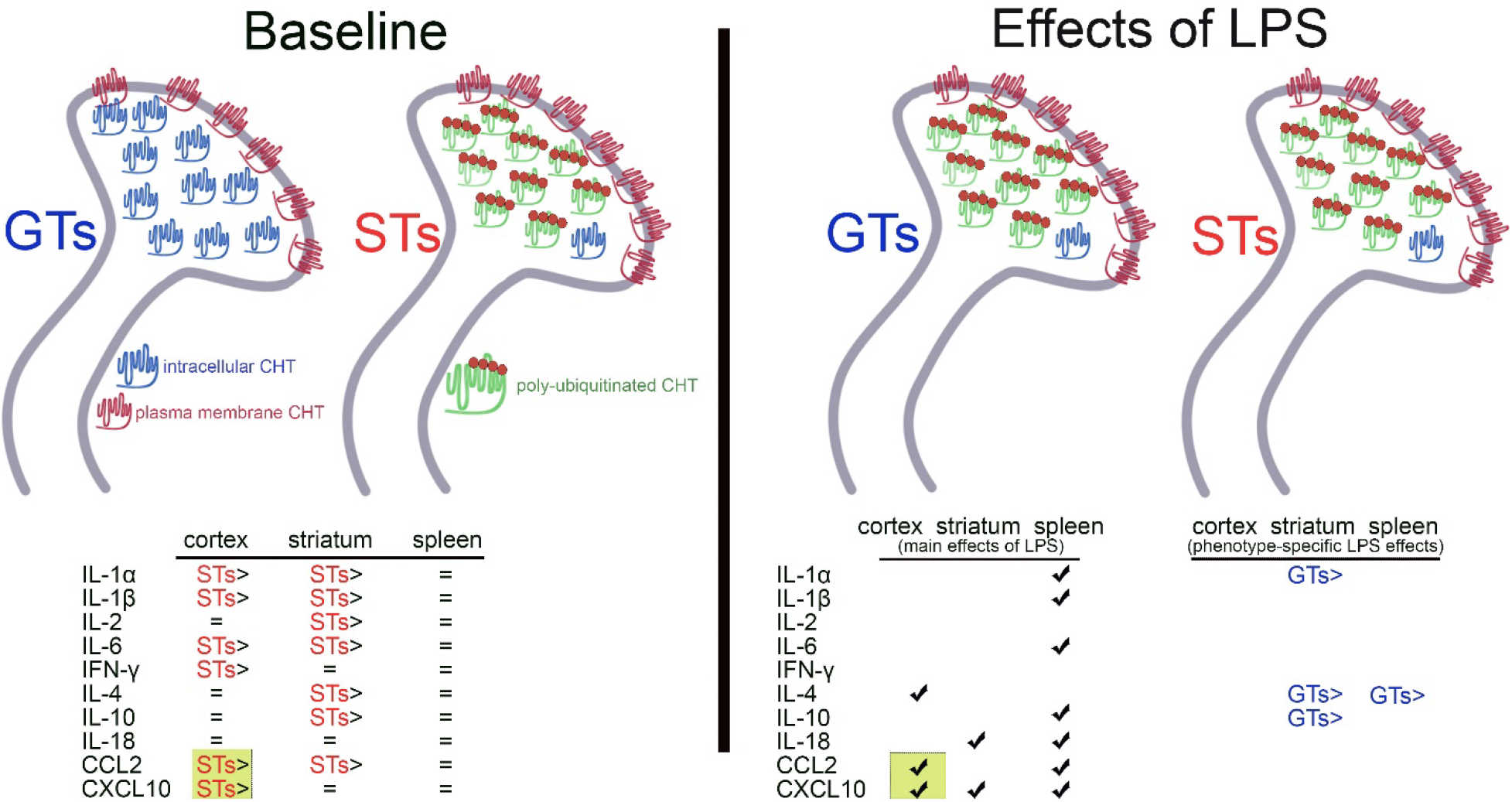
Schematic illustration and summary of results. At baseline (left half of the figure), highly robust increases in levels of poly-ubiquitinated CHTs (green symbols with red dots) were found in STs and subsequently located specifically in vesicular membrane-enriched, intracellular domains. In contrast, in both phenotypes, CHTs located in synaptosomal plasma membrane remained unmarked by ubiquitin (blue symbols), suggesting that they continue to support trans-membrane choline transport. Intracellular ubiquitinated CHTs in STs may be unavailable for outward trafficking, thereby reducing plasma-membrane CHT density, choline transport, and cholinergic activity upon stimulation of cholinergic neurons (Koshy Cherian et al., 2017). At baseline, a range of cytokines and chemokines was upregulated in STs over GTs (bottom left). None of the analytes measured in spleen indicated phenotype-specific differences. Following the treatment with LPS (right half), levels of ubiquitinated CHTs in GTs reached those seen in STs. LPS did not further increase such levels in STs, suggesting a ceiling effect. Furthermore, following LPS treatment, levels of three brain cytokines/chemokines, and 8/10 cytokines from spleen, were elevated in both phenotypes (left part of the bottom table, indicating main effects of LPS). Concerning phenotype-specific effects of LPS (right part of the bottom table, reflecting significant interactions between the effects of phenotype and LPS), levels of 3 brain cytokines and one chemokine (CXCL10), and of IL-4 in spleen, were increased in GTs relative to their baseline levels. In no case did LPS increase cytokine/chemokine levels solely in STs. Only the levels of two analytes and in one brain region - the two chemokines in cortex - were higher in STs at baseline as well as upregulated by LPS (marked by greenish background boxes). Therefore, these two chemokines are most likely to contribute to the potentially complex signaling mechanisms that link neuro-immune modulation with CHT status, and consequently with the cognitive-motivational phenotype indexed by sign-tracking.

The present experiments investigated a new cellular mechanism that potentially contributes to the dampening of CHT function in STs. The cholinergic synaptic capacity limitation that results from such attenuated CHT function has previously been suggested to mediate the attentional control deficits of STs and their associated vulnerability for addiction-like behavior (Koshy Cherian et al., 2017; Pitchers et al., 2018; Pitchers et al., 2017b; Paolone et al., 2013; Phillips and Sarter, 2020). As a first step toward investigating the contributions of a post-translational CHT modification, and the potential causal role of elevated levels of immune modulators for such a modification, the evidence from the present experiments remains in part correlational (in STs, higher levels of intracellular CHT ubiquitination coincided with higher levels of brain cytokine and chemokine levels). The effects of LPS on both measures (elevated ubiquitinated CHTs and brain cytokine levels), however, suggest a causal mechanism. Activation of the innate immune system yielded an increase in CHT ubiquitination, thereby potentially revealing a causal link between activation of neuro-immune modulators and CHT function.

Closer inspection of the present data on the effects of LPS further supports the possibility that elevated brain cytokine levels are part of a causal signaling mechanism via which LPS leads to the ubiquitination of intracellular CHTs. Such an inspection requires first addressing the possibility that ceiling effects in STs prevented the manifestation of further, LPS-induced increases of ubiquitinated CHTs and brain cytokines in this phenotype. Concerning the former, baseline levels in STs (Fig. 2) were nearly at or above ubiquitination levels seen in GTs following LPS administration (Fig. 3). Moreover, administration of a relatively high dose of LPS (5.0 mg/kg) did not further increase CHT ubiquitination levels in GTs seen after the lower dose (1.0 mg/kg), suggesting corresponding maximum concentrations of ubiquitinated CHTs in both phenotypes.

Ceiling effects may have also prevented further LPS-induced increases in cytokine concentrations - except for the two chemokines (below) - in exclusively in STs. The widespread and robust LPS-induced increases in spleen cytokine levels of both STs and GTs indicated the overall efficacy of the endotoxin in both phenotypes. Main effects of LPS on brain cytokine levels, however, were relatively sparse, elevating only IL-4 in cortex and IL-18 in striatum. Importantly, at baseline, neither IL-4 nor IL-18 levels were higher in STs. Thus, none of the brain cytokines that were elevated in STs at baseline were further elevated by LPS (main effects). Moreover, whenever LPS elevated cytokine levels in just one phenotype (indicated by significant interactions between the effects of phenotype and LPS), these increases occurred solely in GTs. These data mirror the effects of LPS on CHT ubiquitination, with baseline cytokine levels in STs already at a maximum concentration.

CCL2 and CXCL10 followed a different pattern. Baseline concentrations of CCL2 and CXCL10 were higher in the cortex, and of CCL2 in striatum, of STs. Yet LPS further increased levels of both in cortex (main effects of LPS) but only of CXCL2 in striatum. Moreover, the magnitude of the LPS-induced increase on CXCL10 levels was noteworthy, reaching nanogram concentrations compared with picogram levels at baseline (Fig. 4). As we hypothesized that levels of brain cytokine and chemokine levels influence CHT ubiquitination at baseline, and that LPS-induced elevation of such levels yields additional CHT ubiquitination, the only two cytokines which met this combined criterion (higher baseline levels in STs and robust LPS-induced increases) were the two chemokines CCL2 and CXCL10 in cortex. Thus, the present results suggest that cortical chemokines are a member of the potentially complex signaling chain that links neuro-immune activation with CHT ubiquitination.

Chemokines orchestrate not just the activation of highly diverse, immune response-supporting signaling pathways but are also involved in numerous cellular regulatory mechanisms not considered part of immune activation and immune surveillance (e.g., Moser et al., 2004). In brain, chemokines are primarily secreted from microglia and neurons, and receptors for chemokines are expressed by neurons and all types of glial cells. Brain chemokines can influence synaptic mechanisms in part via astrocyte-synapse interactions (Biber et al., 2002; Bezzi et al., 2001; Adler and Rogers, 2005; Melik-Parsadaniantz and Rostene, 2008; Sowa and Tokarski, 2021). Thus, elevated of levels of brain chemokines may not necessarily indicate an active neuro-immune response but, independent from their chemotactic capacity, may serve as a neuro-modulatory system that is neuronally regulated and in turn influences neuronal communication and behavior (e.g., Vitkovic et al., 2000; Prieto and Cotman, 2017; Vezzani and Viviani, 2015; Bourgognon and Cavanagh, 2020; Donzis and Tronson, 2014; Davis and Krueger, 2012).

The two chemokines measured in the present studies, CCL2 and CXCL10, are exemplary members of such a neuro-modulatory system. These chemokines and their cognate receptors are expressed in the non-inflamed brain, including in neurons (Banisadr et al., 2002; Mizuno et al., 2003; Xia et al., 2000; Vinet et al., 2010), and they have been shown to affect synaptic, behavioral and cognitive functions (Melik-Parsadaniantz and Rostene, 2008; Nelson et al., 2011; Vlkolinsky et al., 2004). Of particular interest in the present context is the demonstration that the C-C chemokine receptor type 2 (CCR2), the main receptor for CCL2, is expressed by cholinergic neurons in the basal forebrain (Banisadr et al., 2005). Elevated levels of CCL2 in cholinergic neurons therefore may have direct access to regulatory or transcriptional pathways leading to CHT ubiquitination and attenuated cholinergic activity (Silverman et al., 2015). Conversely, as increases in brain cholinergic signaling serve to suppress cytokine levels (Shi et al., 2009; Hao et al., 2011), diminished cholinergic signaling may contribute to heightened cytokine levels (see also Leite et al., 2016), giving rise to bidirectional interactions between the two systems.

In the present experiments, the effects of LPS on CHT ubiquitination and cytokine concentrations served to test the hypothesis that the two measures are mechanistically inter-related, but not to necessarily indicate the possibility that STs express, as a trait component, an activated immune response. This view is strongly supported by the absence of phenotypic differences in spleen cytokine levels. However, as brain levels of several cytokines appeared to be at ceiling levels in STs, at least relative to the efficacy of LPS to further elevate these levels, the notion that brain cytokine levels in STs reflect a considerable degree of neuro-inflammation cannot be dismissed. Additional analyses of brain inflammation markers in STs and GTs were beyond the scope of the present research but will be needed to further explore this possibility.

Although the present LPS effects suggest that brain elevated chemokine levels influence CHT ubiquitination levels, it is important to reiterate that, *vice versa*, attenuated cholinergic signaling promotes brain cytokine expression (references above). Thus, the search for the molecular and genetic causes for such complex, bidirectional interactions may be extremely challenging, despite of the considerable reliability, across laboratories and time, of sign-tracking as a trait. Genome-wide association studies have begun identifying several possible variants associated with sign-tracking; however, these variants differed across rats obtained from individual vendors (Gileta et al., 2022; Fitzpatrick et al., 2013), further complicating such efforts. The present findings may motivate a more directed search for associations with genetic variants which specifically influence brain chemokine expression and the ubiquitination system.

The present results extend the search for the cellular underpinnings of the neuro-behavioral trait indexed by sign-tracking to include the state of the neuro-immune modulator system. Given emerging evidence indicating that drugs of abuse *per se* can activate such neuro-immune modulators (for review see Lacagnina et al., 2017; Crews, 2012), pre-existing elevated brain cytokine concentrations may interact with addictive drug use to facilitate the severity of the neuronal and behavioral characteristics of vulnerability traits, and eventually the manifestation of compulsive drug taking.

## Acknowledgment

We thank Kevin Cordova and Nolan Hamilton (Temple University) for assistance with subcellular fractionation and immunoblotting studies.

